# The single flagellum of *Leishmania* has a fixed polarisation of its asymmetric beat

**DOI:** 10.1101/2020.03.21.001610

**Authors:** Ziyin Wang, Tom Beneke, Eva Gluenz, Richard John Wheeler

## Abstract

Eukaryotic flagella undergo different beat types necessary for their function. The single flagellum on *Leishmania* parasites, for example, undergoes a symmetric tip-to-base beat for forward swimming and an asymmetric base-to-tip beat to rotate the cell. Asymmetric beats are most commonly associated with multi-ciliated tissues or organisms where the asymmetry has a constant polarisation. We asked whether this also holds for the single *Leishmania* flagellum. To do so, we used high frame rate dual colour fluorescence microscopy to visualise intracellular and intraflagellar structure in live swimming cells. This showed that the asymmetric *Leishmania* beat has a fixed polarisation. As in *Chlamydomonas*, this asymmetry arose from an asymmetric static curvature combined with a symmetric dynamic curvature. Some axoneme protein deletion mutants give flagella which retain static curvature, but lack dynamic curvature. We saw that these retain a fixed polarisation. Similarly, deletion mutants which disrupt vital asymmetric extra-axonemal and rootlet-like flagellum-associated structures also retain a fixed polarisation. This indicated that beat asymmetry does not originate from rootlet-like and extra-axonemal structures and is likely intrinsic to either the nine-fold rotational symmetry of the axoneme structure or due to differences between the outer doublet decorations.

## Introduction

Motile flagella and cilia have essentially indistinguishable ultrastructures but originally received different names based on their biological function – a combination of where they are present in organisms, their structure and the motion they undergo (Takeda and Narita, 2012). The organelle tends to be called a flagellum when they undergo a symmetric planar near-sinusoidal beat and there are few per cell. In contrast, the term cilium tends to be used when they undergo an asymmetric planar wafting beat and there are many cilia per cell or many ciliated cells undergoing coordinated movement across a tissue. Ultimately, the correct choice of symmetric or asymmetric waveform, and the correct polarisation of the latter, must be used to achieve the necessary biological function. Defects tend to cause motility defects in swimming cells and ciliopathies, a range of mild to severe genetic diseases, in humans.

Typical asymmetric beats undergo a power stroke, which drives fluid movement relative to the cell, followed by a recovery stroke, returning the cilium/flagellum to its starting configuration. A planar asymmetric beat has two possible polarisations, corresponding to which way the power stroke pushes fluid as the flagellum beats. In multi-ciliated systems the power stroke’s polarisation tends to be organised to generate fluid flows (e.g. animal ciliated epithelia, such as in brain ventricles (Faubel et al., 2016)) or to drive cell swimming (e.g. *Tetrahymena* and *Paramecium*) (Naitoh and Kaneko, 1972). There are varied configurations in cells with fewer cilia/flagella, for example *Chlamydomonas* uses asymmetric beating of two flagella with opposite polarisation to achieve forward swimming (Ringo, 1967) while the unicellular eukaryotic parasite *Leishmania* uses asymmetric beats of its single flagellum to rotate (Gadelha et al., 2007; Holwill and McGregor, 1975) (Figure 1A,B). Cilia/flagella have asymmetries which contribute to generating the correct beat form – firstly those which keep bending in a plane, and secondly those which introduce asymmetries in the beat with the correct polarisation. Although both structural and functional (e.g. signalling) asymmetries may contribute, far more is known about structural asymmetries.

**Figure 1.**
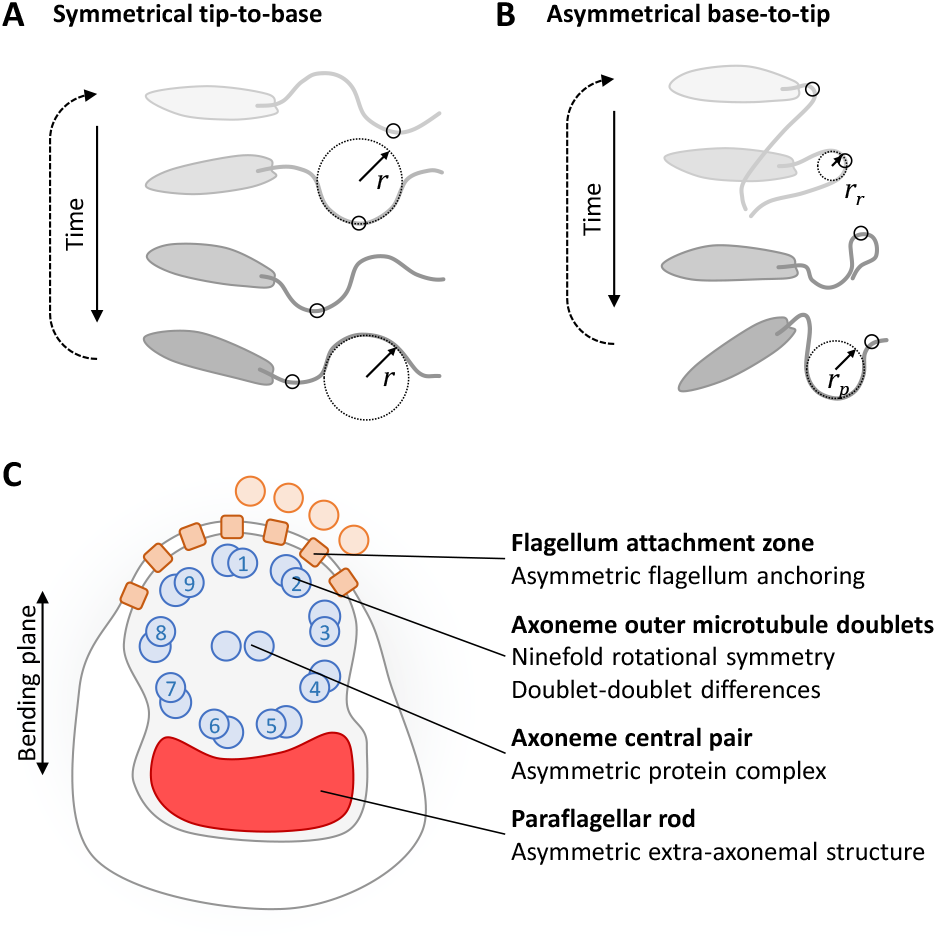
Known asymmetries in *Leishmania* flagellar beating and flagellum ultrastructure. **A-B.** Cartoon representations of the two types of *Leishmania* beat. **A.** The symmetric beat where waves propagate from the tip to the base with symmetric curvature (r). **B.** The asymmetric beat where waves propagate from the base to the tip with asymmetric curvature for power and recovery strokes (r_p_ and r_r_) **C.** Cartoon representation of key asymmetries in axoneme structure which may contribute to asymmetry of the base-to-tip beat. This includes asymmetries in the central pair and outer doublets of the axoneme, the paraflagellar rod and (toward the flagellum base) the attachment to the cell body by the flagellum attachment zone.

Structural asymmetries for keeping bending in a plane are well-characterised. Motile flagella/cilia canonically have nine outer microtubule doublets in a circular arrangement, around a central pair of singlet microtubules (a 9+2 arrangement). Their beating is driven by coordinated activity of dynein motors bound to the outer doublet microtubules, which drive sliding between adjacent doublets. Flagella with nine-fold rotational symmetry of the outer doublets and no central pair or asymmetric extra-axonemal structures tend to undergo three-dimensional rotating or helical movement, such as nodal cilia (Nonaka et al., 1998). To achieve a planar beat, this rotational symmetry must be broken. Depending on the organism this may involve addition of a fixed orientation central pair complex (Ishijima et al., 1988), which does not have reflectional or rotational symmetry (Carbajal-González et al., 2013). Alternatively, it may involve the presence of specialised bridges between particular microtubule doublets to restrict where dynein-driven sliding occurs (Bui et al., 2009; Gibbons, 1961) thus removing the nine-fold rotational symmetry of the outer doublets.

The nature of structural asymmetries which a) give rise to asymmetric beats and b) defines their polarisation are less well understood, however many asymmetries are known which correlate with asymmetric beat polarisation. Within the 9+2 axoneme, differences between the outer doublet decoration, particularly the inner dynein arms, are likely important (Bui et al., 2009; Bui et al., 2012). In addition to the asymmetric 9+2 axoneme, cilium-associated structures tend to be asymmetric. This includes the anchoring of the basal body to the cell by rootlet structures – such as the basal body-basal body linkage by the distal striated fibre in *Chlamydomonas* (Dutcher and O’Toole, 2016; Ringo, 1967), the basal foot structure in animal ciliated epithelia cells (Gibbons, 1961) and the basal body-associated structures in *Paramecium* (Tassin et al., 2016). Mutations of rootlet proteins cause a loss of ciliated tissue polarity. However, individual cilia rotate randomly while retaining their normal asymmetric structure (Clare et al., 2014; Kunimoto et al., 2012). Each individual cilium presumably retains their asymmetric beat, although this has not been analysed in detail.

An asymmetric beat can arise when a flagellum has a large static curvature (underlying ‘shape’ of the flagellum) in addition to the symmetric dynamic curvature (the propagating wave) (Eshel and Brokaw, 1987; Geyer et al., 2016). In *Chlamydomonas*, several mutants are known which have a more symmetric beat, including *pf2* (Brokaw and Kamiya, 1987) and *mbo2* (Segal et al., 1984). These mutants have a greatly reduced static curvature (Geyer et al., 2016). However, no known mutants invert the bend of this static component to invert the asymmetry of the beat.

It is unknown whether a single *Leishmania* flagellum has a fixed polarisation for its asymmetric beat. We therefore asked whether polarisation is fixed or switchable and what structures could be involved (Figure 1C). In addition to the 9+2 axoneme, *Leishmania* have an extra-axonemal structure called the paraflagellar rod (PFR). There are several possible functions of the PFR (Portman and Gull, 2010) which is required for normal flagellum beating (Santrich et al., 1997). Its protein composition indicates Ca^2+^ and cAMP regulation roles (Oberholzer et al., 2007; Portman et al., 2009), while its structure suggests biomechanical effects to convert planar to three dimensional movement (Hughes et al., 2012; Koyfman et al., 2011) although we have previously argued against the latter (Wheeler, 2017). The PFR has a fixed asymmetric position and is attached to outer microtubule doublets 4-6 (Fuge, 1969; Gadelha et al., 2005). The anchoring of the flagellum base to the cell is also asymmetric, analogous to rootlet structures in other organisms, with lateral attachment to the cell body in the flagellar pocket via the flagellum attachment zone (FAZ) only near outer microtubule doublets 9, 1 and 2 (Wheeler et al., 2016). The 9+2 axoneme itself likely also has asymmetries between outer microtubule doublets, based on recent cryo-electron tomography of the related species *Trypanosoma brucei* (Imhof et al., 2019).

Analysing polarisation of *Leishmania* asymmetric beats has a key challenge: *Leishmania* cells appear axially symmetric by transmitted light illumination methods. To determine the cellular orientation in swimming cells we developed high frame rate dual colour widefield epifluorescence of *Leishmania* promastigotes to observe an asymmetric internal cytoskeletal structure (labelled with a fluorescent protein) while also observing flagellum beating as the cell undergoes different flagellum beat behaviours. By combining this with mutations in different asymmetric features of the flagellum and cell-flagellum attachment we showed 1) that the flagellum has a fixed polarisation of the asymmetric flagellar beat, 2) paralysed flagellum mutants which form a static curvature retain a fixed polarisation and 3) this asymmetry does not require the large PFR structure or lateral attachment by the FAZ. This has implications for the mechanisms by which this parasite may achieve directed taxis.

## Results

*Leishmania* flagellum beating can be readily analysed from high frame rate (400 Hz) phase contrast videos using automated tracing of the flagellum configuration (Walker and Wheeler, 2019; Walker et al., 2019). As often used for other organisms, this data can be represented as plots of tangent angle at different distances along the flagellum over time (Figure 2, Video 1), where tangent angle refers to the angle of a section of flagellum relative to the flagellum base. *Leishmania* undergo two well-described types of flagellum beating, a high frequency tip-to-base beat or a lower frequency base-to-tip beat. These occur at around 20 to 25 Hz for the tip-to-base and around 5 Hz for the base-to-tip beats (Gadelha et al., 2007; Wheeler, 2017). The tip-to-base beat is symmetric (typical of flagella) while the base-to-tip beat is asymmetric with a power and recovery stroke (typical of cilia) (Gadelha et al., 2007; Holwill and McGregor, 1975).

**Figure 2.**
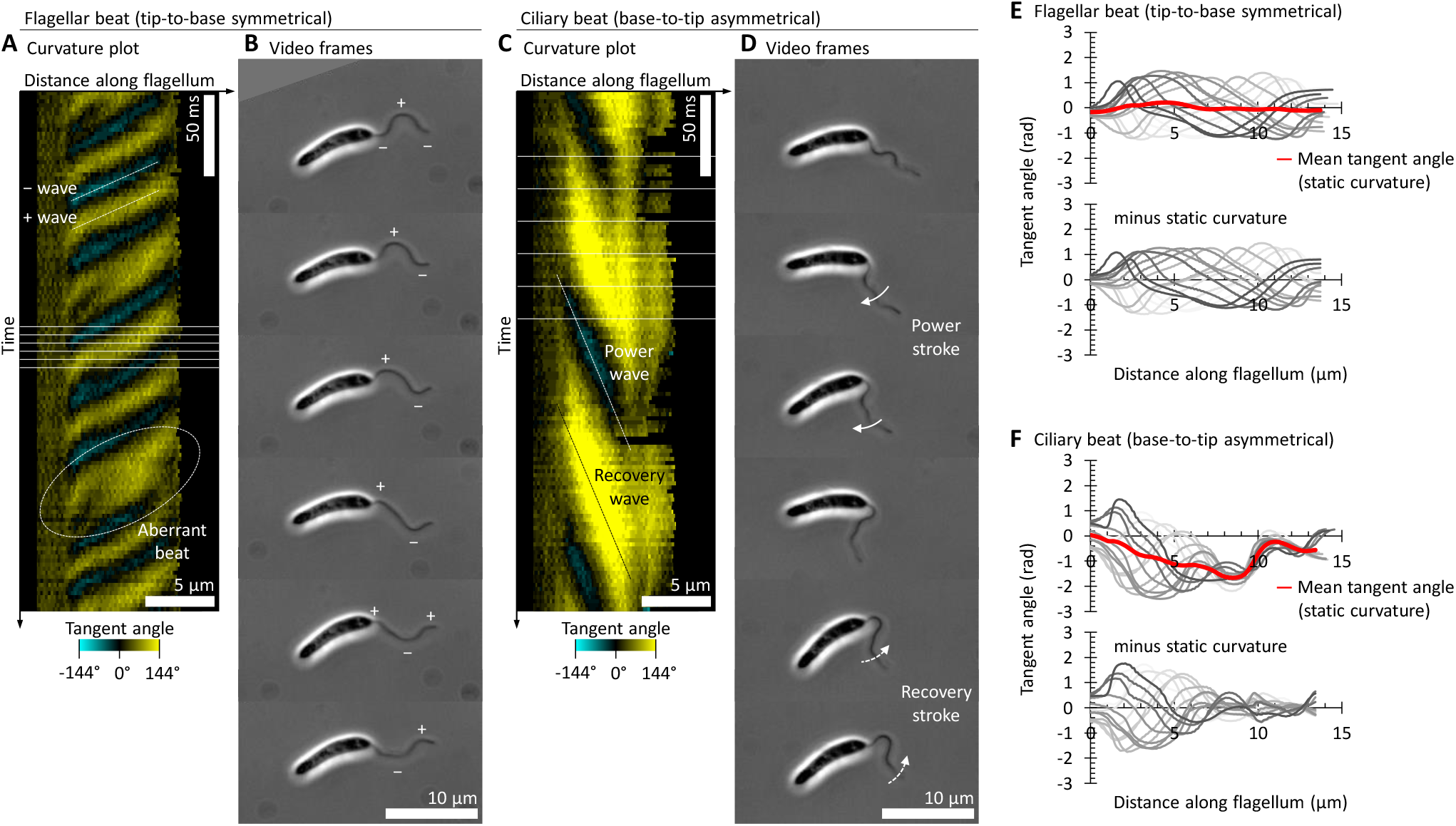
The symmetric and asymmetric beats of the *Leishmania* flagellum. One example cell undergoing either **A-B.** a symmetric tip-to-base flagellar beat or **C-D.** an asymmetric base-to-tip ciliary beat visualised from a 400 Hz high-frame rate phase contrast video. **A.** Flagellum curvature over time for the symmetric beat, plotted as a kymograph of tangent angle along the flagellum, automatically traced from the phase contrast video. Yellow indicates clockwise tangent angles relative to the flagellum base and cyan anticlockwise. The waves during the symmetric beat propagate from the flagellum tip to its base (from top right to bottom left) alternating between waves with a negative tangent angle (labelled − wave) or a positive tangent angle (labelled + wave). Aberrant waves occasionally occur with a different frequency or with additional bends in the flagellum, one example is indicated. **B.** Six frames illustrating the propagation of positive and negative tangent angle waves over one beat cycle. Phase contrast images from the frames indicated with horizontal lines in A. **C.** Flagellum curvature over time for the asymmetric beat. The waves propagate from the flagellum base to its tip (from top left to bottom right) alternating between a wave for a power stroke (small tangent angles, negative in this example) and a wave for a recovery stroke (large tangent angles, positive in this example). **D.** Six frames illustrating the power and recovery strokes over one beat cycle. Phase contrast images from the frames indicated with horizontal lines in C. **E.** Top, plotted tangent angles at different distances along the flagellum for one symmetric tip-to-base beat cycle in **A-B**. Mean tangent angle at each distance along the flagellum gives the static curvature of the flagellum, which is very near straight. Bottom, tangent angles with static curvature subtracted. **F.** Plotted tangent angle as in **E**, except for one asymmetric base-to-tip beat cycle in **C-D**. Mean tangent angle follows a negative approximately linear trend in the proximal flagellum. Subtraction of the static curvature shows a near-symmetric dynamic component, with decreasing amplitude from the proximal to distal flagellum.

Using tangent angle plots the wavefronts of the high frequency tip-to-base symmetric beat appear as lines aligned top right to bottom left, alternately with positive or negative tangent angles (Figure 2A, Video 1A). These correspond to the positive and negative curvature wavefronts propagating from the flagellum tip over time (Figure 2B). The wavefronts of the asymmetric base-to-tip beat appears as wider diagonal lines aligned top left to bottom right corresponding to lower frequency waves propagating from the flagellum base over time (Figure 2C, Video 1B). Unlike the symmetric beat, the alternating positive and negative tangent angle wavefronts in the asymmetric beat have different magnitudes of tangent angle – the wavefront for the recovery stroke has tighter curvature giving larger tangent angles than the wavefront for the power stroke (Figure 2C,D).

The static curvature of the flagellum can be determined by averaging the tangent angle over an integer number of beats or by taking the static/infinite frequency mode of a Fourier decomposition. While undergoing a symmetric tip-to-base beat the static curvature is typically near-zero (Figure 2E). In contrast, for asymmetric base-to-tip beats the static curvature reaches large values (>*π*/2 rad, >90°) (Figure 2F). The dynamic curvature of the flagellum can be determined by subtracting the static curvature. This corresponds to the propagating wave shape on the underlying static shape of the flagellum. The dynamic curvature of both the tip-to-base and base-to-tip beats are near-symmetric, however the base-to-tip beats often do not propagate along the entire flagellum (Figure 2F). For the portions of the flagellum with a large-amplitude base-to-tip beat the static curvature is near-linear, similar to observed in *Chlamydomonas* asymmetric beats (Geyer et al., 2016).

### The asymmetric *Leishmania* beat occurs with a constant polarisation

*Leishmania* have a fixed axoneme central pair orientation (Figure 1C)(Gadelha et al., 2006) and bending for flagellum beating is thought to occur only in the plane perpendicular to the central pair. Therefore, there are two possible directions for the power stroke of the asymmetric beat. Ciliary beating in other organisms is highly polarised, suggesting a single preferred direction, however the *Leishmania* cell is near-axially symmetric meaning cell orientation is not visible from transmitted light images like phase contrast (Figure 2). Whether there is a preferred direction can be tested by observing asymmetric intracellular structures, which act as a reporter of cell orientation, during flagellum beating. We achieved this using dual colour high frame rate widefield epifluorescence microscopy (Figure S1) using a cell line expressing a well-characterised flagellum membrane marker, SMP1::mCh (Tull et al., 2004; Wheeler et al., 2015), and a marker of the asymmetric microtubule-based cytoskeletal structure comprising the microtubule quartet and the lysosomal microtubule(s), mNG::SPEF1 (Gheiratmand et al., 2013; Halliday et al., 2018; Wang et al., 2019) (Figure 3A).

**Figure 3.**
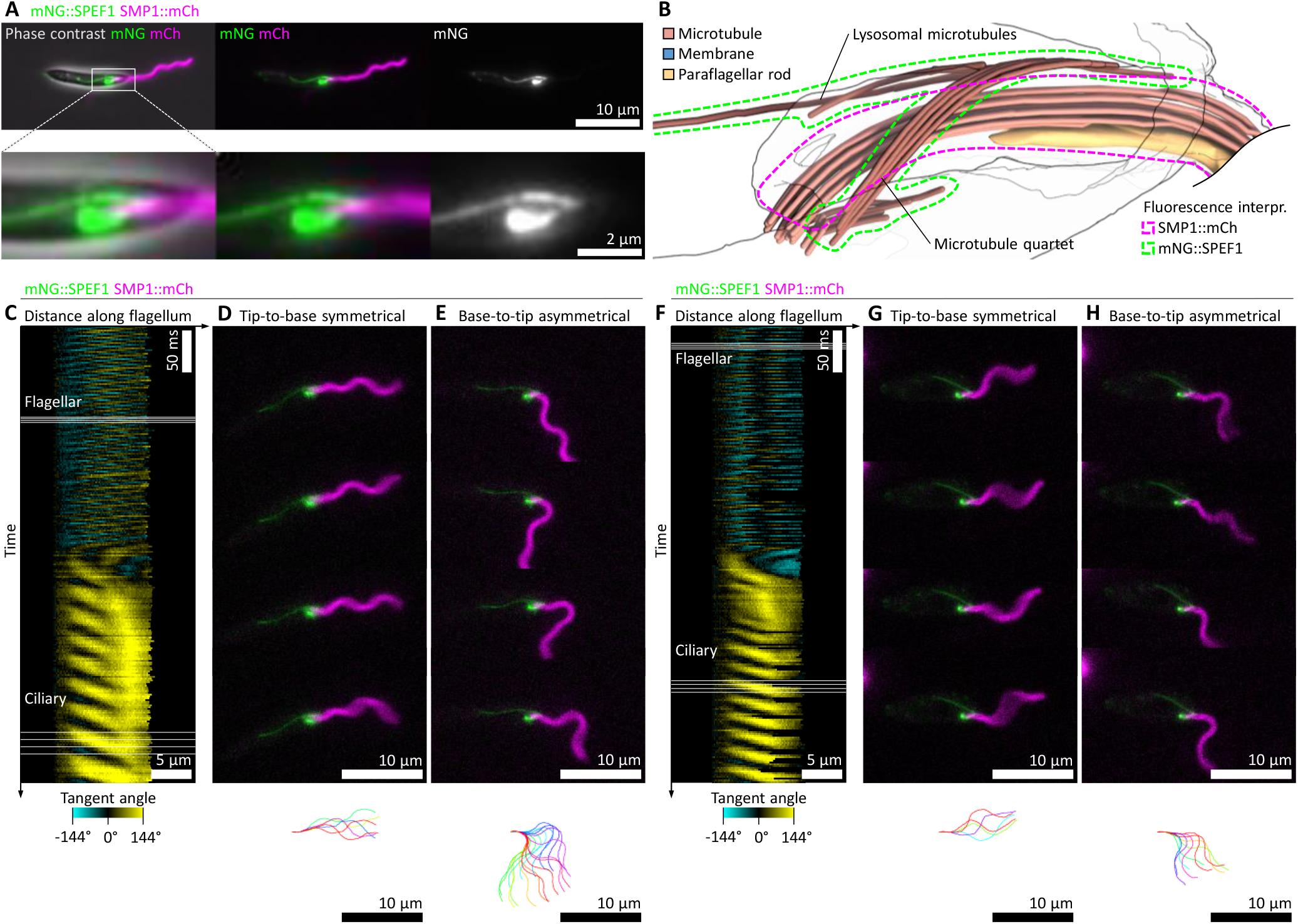
SPEF1 is an asymmetric reporter of *Leishmania* cell orientation during asymmetric flagellum beating. **A.** The subcellular localisation of the microtubule quartet and lysosomal microtubule-associated mNG::SPEF1 relative to the flagellar membrane marker SMP1::mCh by epifluorescence microscopy. **B.** Structure of the flagellar base and flagellar pocket by electron tomography, as previously published (Wheeler et al., 2016), indicating the SPEF1- and SMP1-associated ultrastructures. **C-H.** Two examples of flagellum movement and cell orientation in cells expressing mNG::SPEF1 and SMP1::mCh switching from a symmetric tip-to-base to an asymmetric base-top-tip beat, derived from 100 Hz high-frame rate dual colour epifluorescence videos. **C-E.** First example cell. **F-H.** Second example cell. **C,F.** Flagellum curvature over time, plotted as a kymograph of tangent angle along the flagellum, automatically traced from SMP1::mCh signal. Yellow indicates tangent angles to the right, cyan to the left. **D,G.** Four frames showing the symmetric tip-to-base beat. mNG::SPEF1 and SMP1::mCh fluorescence from the frames indicated “Flagellar” in C,F. The flagellum configuration over one beat cycle is shown at the bottom. **E,H.** Four frames showing the asymmetric base-to-tip beat. mNG::SPEF1 and SMP1::mCh fluorescence from the frames indicated “Ciliary” in C,F. The asymmetric beat tends to bend away from the side of the cell with the lysosomal microtubule.

Through comparison to previously published electron tomography of the flagellar pocket (Wheeler et al., 2016) the relative orientation of the flagellum axoneme, paraflagellar rod and flagellum attachment zone can be inferred from the appearance of the mNG::SPEF1 signal (Figure 3B). In Figure 3B, the plane of beating is the plane of the image (perpendicular to the central pair). As the flagellum exits the cell through the flagellar pocket neck it attaches to the microtubule quartet via the flagellum attachment zone on one side of the flagellum. This attachment region is near where specialised lysosome-associated microtubule(s) also meet the microtubule quartet, on the opposite side of the flagellum to the start of the paraflagellar rod (Wheeler et al., 2016).

In high frame rate videos the flagellum movement can be visualised and traced from the SMP1::mCh signal. Identifying example cells where the beat switches from symmetric to asymmetric over the course of the video is relatively easy, illustrated for two example cells in Figure 3C-H & Video 2. The switch between flagellum beats is summarised by tangent angle at different distances along the flagellum over time (Figure 3C, F). In these videos, the orientation of bending relative to the cellular ultrastructure can be inferred from the mNG::SPEF1 signal, both while undergoing a symmetric beat (example frames and the traced beat shown in Figure 3D,G) or an asymmetric beat (Figure 3E,H). This showed the power stroke of the asymmetric beat pushing away from the side of the cell with the lysosomal microtubule(s) corresponding to a static curvature toward that side of the cell.

We noted that the symmetric to asymmetric beat switching involved 1) a tip-to-base wave stalls then reverses, giving a bend which gradually propagates towards the flagellum tip. 2) High frequency tip-to-base waves continue to initiate near the flagellum tip but fail to propagate past the stalled/reversed bend. 3) Lower frequency asymmetric base-to-tip waves start to initiate at the flagellum base but fail to propagate past the stalled/reversed bend. This can give both base-to-tip waves in the proximal domain and tip-to-base waves in the distal domain of a single flagellum at the same time (which is unlike previously described (Gadelha et al., 2007)). 4) Finally, base-to-tip waves can propagate along the whole flagellum and no new tip-to-base waves initiate. The gradually propagating bend in 1) is likely the base-to-tip establishment of the static curvature for the asymmetric beat.

### The paraflagellar rod is on the inside of the tightly curved recovery stroke

The analysis of cell asymmetry using mNG::SPEF1 (Figure 3C-H) in combination with the ultrastructure of the cell (Figure 3B) indicates that the paraflagellar rod sits on the leading side of the flagellum during the asymmetric beat power stroke which corresponds to the inside of the static curvature. To directly confirm this result we analysed two cell lines, one expressing two axoneme markers (mNG::PF16 and mCh::RSP4/6) and one expressing a paraflagellar rod marker and an axoneme marker (mNG::PFR2 and mCh::RSP4/6). In the former the red and green signal should precisely co-localise, while in the latter there should be a small offset – approximately 150 nm based on flagellum ultrastructure (Figure 1C). Again, it was relatively easy to find examples of cells where the beat switches from symmetric to asymmetric (Figure 4, Video 3). The mCh::RSP4/6 signal was much weaker than SMP1::mCh, therefore to analyse these videos we used automated tracing to determine the configuration of the flagellum, then digitally straightened the flagellum from each frame of the video. Multiple frames can then be averaged, to increase signal relative to background noise, allowing precise measurement of red/green signal offset. This showed that there was a precise co-localisation of the two axoneme markers (mNG::PF16 and mCh::RSP4/6. Figure 4A-E, Video 3A), both in straightened images from symmetric beats (Figure 4D) and asymmetric beats (Figure 4E). In contrast the paraflagellar rod was consistently offset from the axoneme (mNG::PFR2 and mCh::RSP4/6, Figure 4F-J, Video 3B). The direction of the offset indicates the paraflagellar rod is on the inside of the static curvature, and the offset was consistent with the expected position of the paraflagellar rod relative to the axoneme.

**Figure 4.**
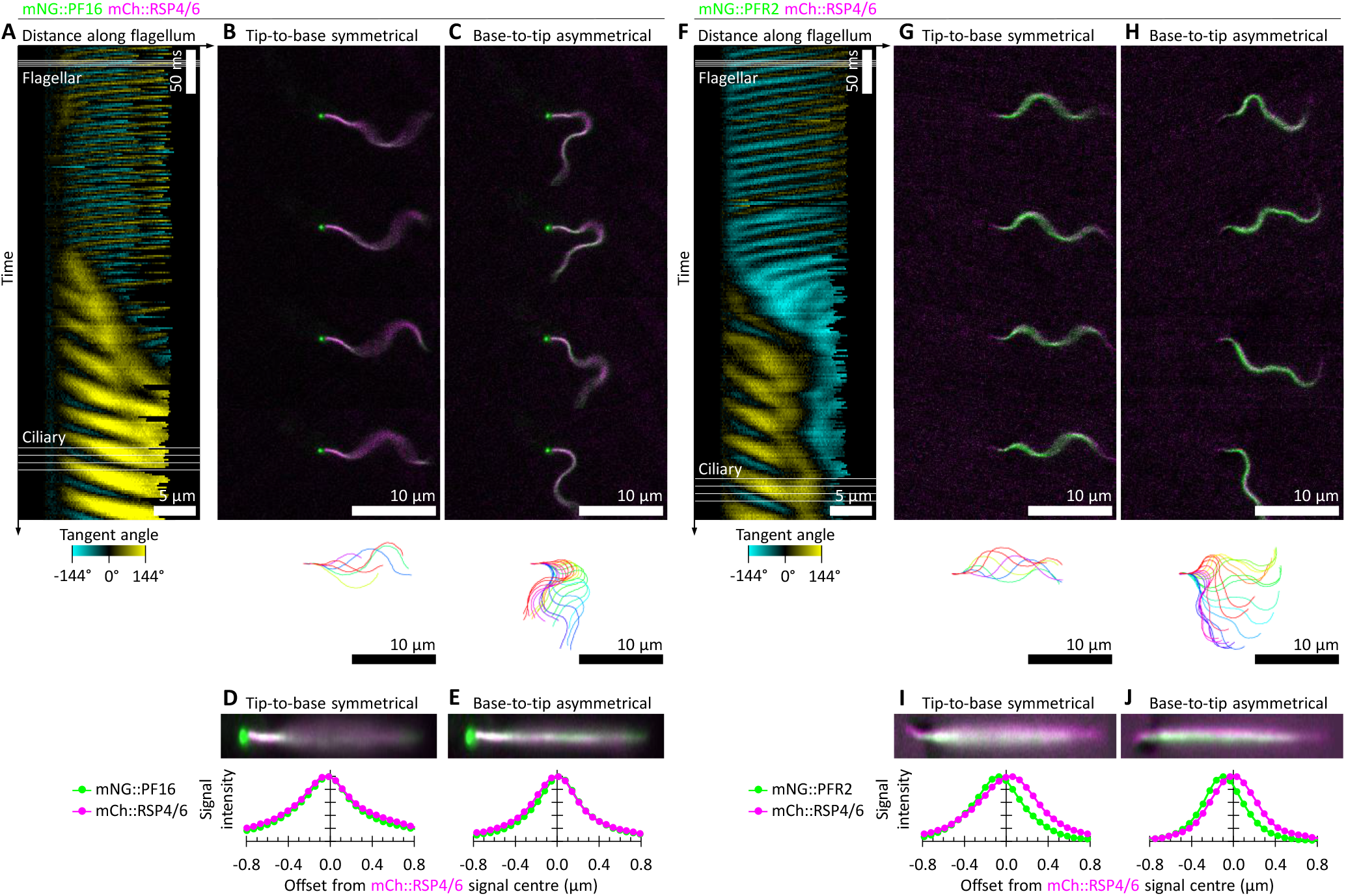
The position of the asymmetric paraflagellar rod extra-axonemal structure in beating flagella. **A-E.** Flagellum movement in a cell expressing mNG::PF16 and mCh::RSP4/6 (two axonemal proteins –central pair and radial spokes respectively) switching from a symmetric tip-to-base to an asymmetric base-top-tip beat, derived from a 100 Hz high-frame rate dual colour epifluorescence video. **A.** Flagellum curvature over time, as in Figure 3C, automatically traced from mNG::PF16 signal. **B.** Four frames showing the symmetric tip-to-base beat and the flagellum configuration over the beat cycle. **C.** Four frames showing the asymmetric base-to-tip beat and the flagellum configuration over the beat cycle. **D.** lly straightened mNG::PF16 and mCh::RSP4/6 signal, averaged over 64 frames of tip-to-base symmetric beating, presented stretched in the transverse direction by 2**×**, and the transverse signal intensity profile. **E.** As for D, but for 64 frames of tip-to-base asymmetric beating. There is no offset between mNG::PF16 and mCh::RSP4/6 signal. **F-J.** As for A-E, except a cell expressing mNG::PFR2 and mCh::RSP4/6 (a paraflagellar rod and an axonemal protein respectively). mNG::PFR2 signal is offset from mCh::RSP4/6 signal. During asymmetric beating the flagellum curves with the paraflagellar rod on the inside of the narrower curvature recovery stroke.

Using many dual colour high frame rate widefield epifluorescence microscopy videos of either the mNG::SPEF1/SMP1::mCh (*n* = 9) or mNG::PFR/mCh::RSP4/6 (*n* = 10) cell lines we can now determine the incidence of the two possible polarisations of the asymmetric beat – either with the power stroke bending away from the side of the cell with the lysosomal microtubules with the paraflagellar rod sitting on the leading side of the flagellum or the inverse (Figure 5A). This indicated a strong and consistent polarisation of the asymmetric beat (Figure 5B), with a tighter radius of curvature of the reverse bend (doublet 1 on the outside of the curve) than the principal bend (doublet 1 on the inside). This indicates the PFR experiences greater compression during the recovery stroke/reverse bend than extension during the power stroke/principal bend.

**Figure 5.**
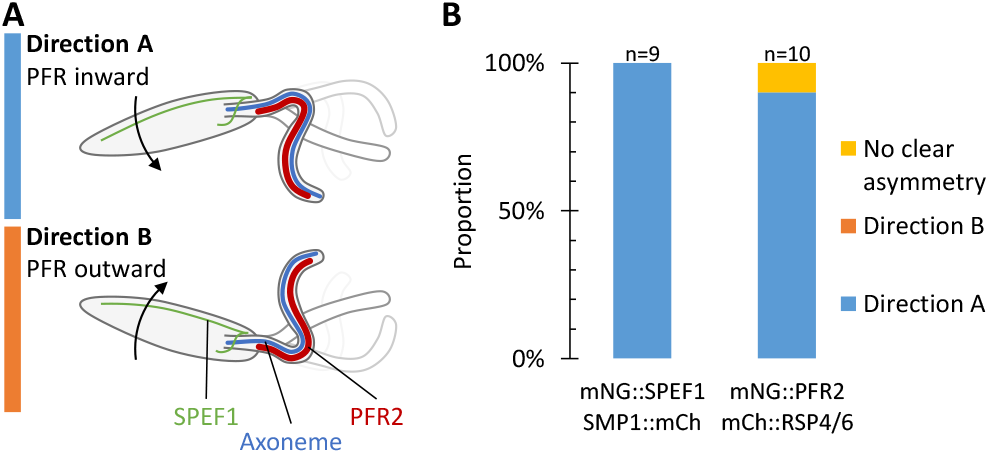
Asymmetric base-to-tip beating occurs in a consistent orientation. **A.** *Leishmania* flagellum beating is near-planar, giving two possible orientations for the asymmetric base-to-tip flagellum beat. Either A, with the PFR on the inside of the narrow radius recovery stroke bend, or B, with the PFR on the outside of the recovery stroke bend. **B.** Instances of asymmetric bending in the A or B direction, counted from high frame rate fluorescence videos of cells expressing mNG::SPEF1 and SMP1::mCh (as in Figure 3) or mNG::PFR2 and mCh::RSP4/6 (as in Figure 4F-J). The asymmetric beat has a strong preference for the power stroke to bend away from the side of the cell with the lysosomal microtubule, positioning the paraflagellar rod on the inside of the tightly curved recovery stroke.

To determine how the asymmetric structures of the *Leishmania* flagellum and associated structures (Figure 1C) contribute to this asymmetric behaviour we analysed flagellum bending in mutants of the axoneme components required for motility, paraflagellar rod, and the flagellum attachment zone.

### Mutants only able to form a static bend have inverted polarisation to asymmetric beats

In *Leishmania* deletion mutants of many conserved axoneme proteins have a paralysed flagellum unable to undergo a beat (lack the dynamic bending component). However in a subset the paralysed flagella retain some capacity for bending and a large proportion have a curled configuration – typically a few turns of a coil(Beneke et al., 2019) – these mutants can form a static but not a dynamic flagellar beat component. We tested whether the static curvature of these mutants retain a preferred polarisation using two mutants where curling is highly prevalent (Beneke et al., 2019). The first was a deletion of the inner dynein arm intermediate chain protein IC140 (Hendrickson et al., 2013; Heuser et al., 2012). The second was a deletion of Hydin, a central pair complex protein required for central pair microtubule stability and to prevent central pair rotation (Dawe et al., 2007). Again, we used mNG::SPEF1 as a reporter to allow determination of the direction of flagellum bending. Curled flagella in both the ΔIC140 and ΔHydin mutant have a preferred bend direction towards the side of the cell with the lysosomal microtubule with the paraflagellar rod on the outside of the coil, although curling in both directions did occur (Figure 6A-D). To confirm this result, we used scanning electron microscopy (SEM) of detergent-extracted cytoskeletons of these two mutants. This allowed direct visualisation of the PFR on the outside face of a large majority of coiled flagella (Figure 6E-H).

**Figure 6.**
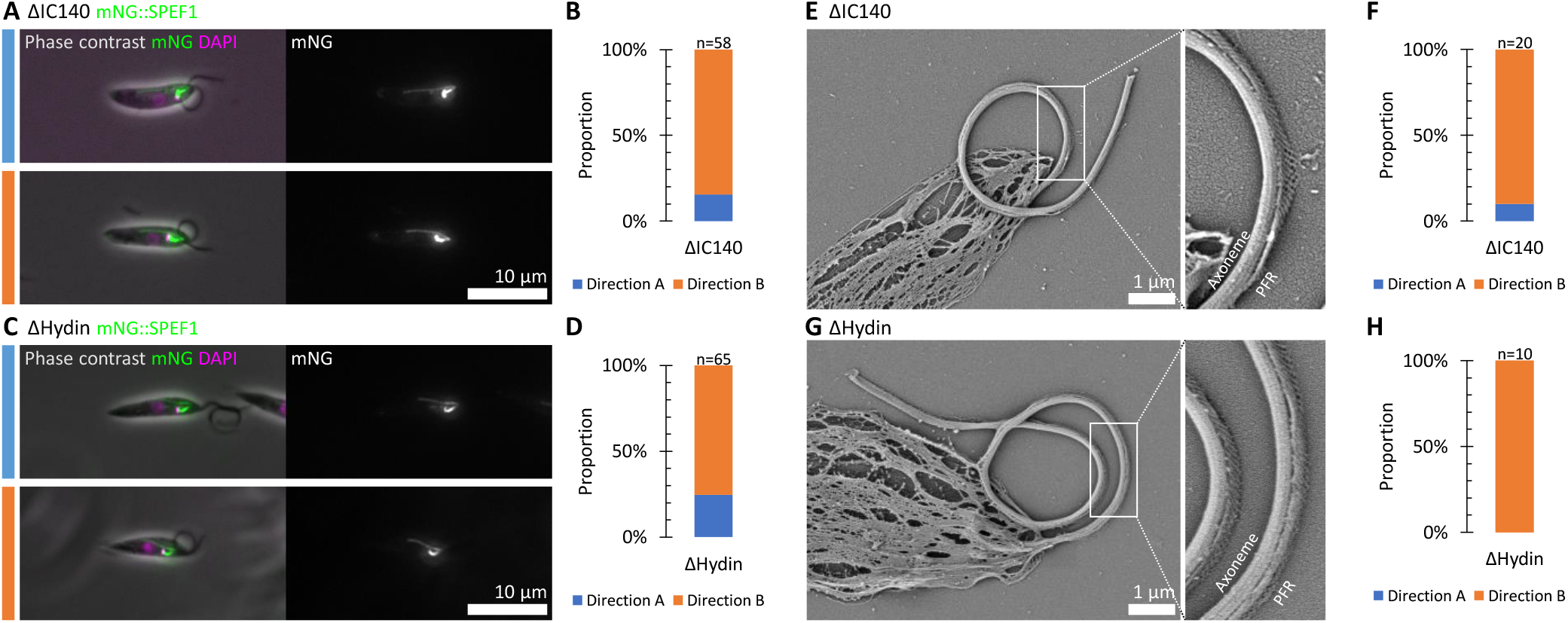
Axoneme protein deletion mutants with a paralysed ‘curled flagellum’ phenotype have a preferred curl direction. **A.** Example fluorescence micrographs of a IC140, an inner dynein arm intermediate chain, deletion mutant expressing mNG::SPEF1, with the flagellum either curling toward the side of the cell with the lysosome microtubule (direction A, paraflagellar rod on the outside of the curve) or the opposite (direction B). **B.** Proportion of cells, assessed from light microscopy, with the flagellum curling in direction A or B. **C-D.** As for A-B, except for a deletion mutant of Hydin, a central pair complex protein. **E.** Example scanning electron micrograph of a detergent-extracted cytoskeleton from an IC140 deletion mutant cell. The direction of curling is directly visible from the PFR next to the axoneme. **F.** Proportion of cells, assessed from scanning electron microscopy, with the flagellum curling in direction A or B. **G-H.** As for E-F, except for a deletion mutant of Hydin. These mutants with dysregulation of flagellum beating still have a preferred bend direction, albeit for a non-beating flagellum.

These curled flagella correspond to a strong static curvature in the opposite direction to the static curvature of the asymmetric base-to-tip beat. The curvature was also much larger, commonly reaching tangent angles of >2*π* rad, >360° (Figure 6A, C). Interpreting this result is difficult as it is not clear that the flagellum curling direction in these mutants originates from the same molecular mechanism of the asymmetry of the normal asymmetric beat. However, it is clear that upon disruption of either the inner dynein arms (ΔIC140) or the central pair (ΔHydin) the flagellum retains the ability for polarised static curvature while losing the ability for dynamic curvature from dysfunction within the 9+2 axoneme.

### Disruption of asymmetric extra-axonemal flagellum structures does not alter polarisation

The paraflagellar rod is a large extra-axonemal structure of comparable size to the axoneme. It is specific to the euglenid lineage of unicellular eukaryotes and is asymmetrically positioned in the flagellum next to doublets 4 to 6 of the axoneme (Figure 5C). PFR2 is a major structural component of the paraflagellar rod and deletion of PFR2 leads to loss of almost all of the paraflagellar rod. In *Leishmania* this leads to a flagellar beat which is still dominantly tip-to-base but with a shorter wavelength and lower amplitude, leading to slower forward swimming (Santrich et al., 1997). A similar motility defect also occurs on PFR2 deletion in the related parasite *Trypanosoma brucei* (Bastin et al., 1999). To determine whether loss of the paraflagellar rod alters the base-to-tip asymmetric beat we generated a PFR2 deletion cell line expressing mNG::SPEF1 and SMP1::mCh (Figure 7, Video 4). Plots of tangent angle over time show the multiple defects the flagellum beat experiences in the absence of the PFR (Figure 7A,D). As previously described, flagella still predominantly undergo base-to-tip waveforms but examples of tip-to-base waveforms were also readily identifiable. Tangent plots confirm that bending tends to be lower amplitude and many wavefronts fail to propagate along the entire length of the flagellum. The frequency was also variable. The lower amplitude flagellum movement also allows the cell to rotate more readily between the slide and coverslip, complicating analysis. Nonetheless, the tip-to-base beat still tends to be near-symmetrical, if a little uncoordinated (Figure 7B, E). The more infrequent base-to-tip beat is still often asymmetric with normal polarity (Figure 7C), however in comparison to the parental line (Figure 3,5) there were also more cells with less pronounced asymmetry (Figure 7F,I).

**Figure 7.**
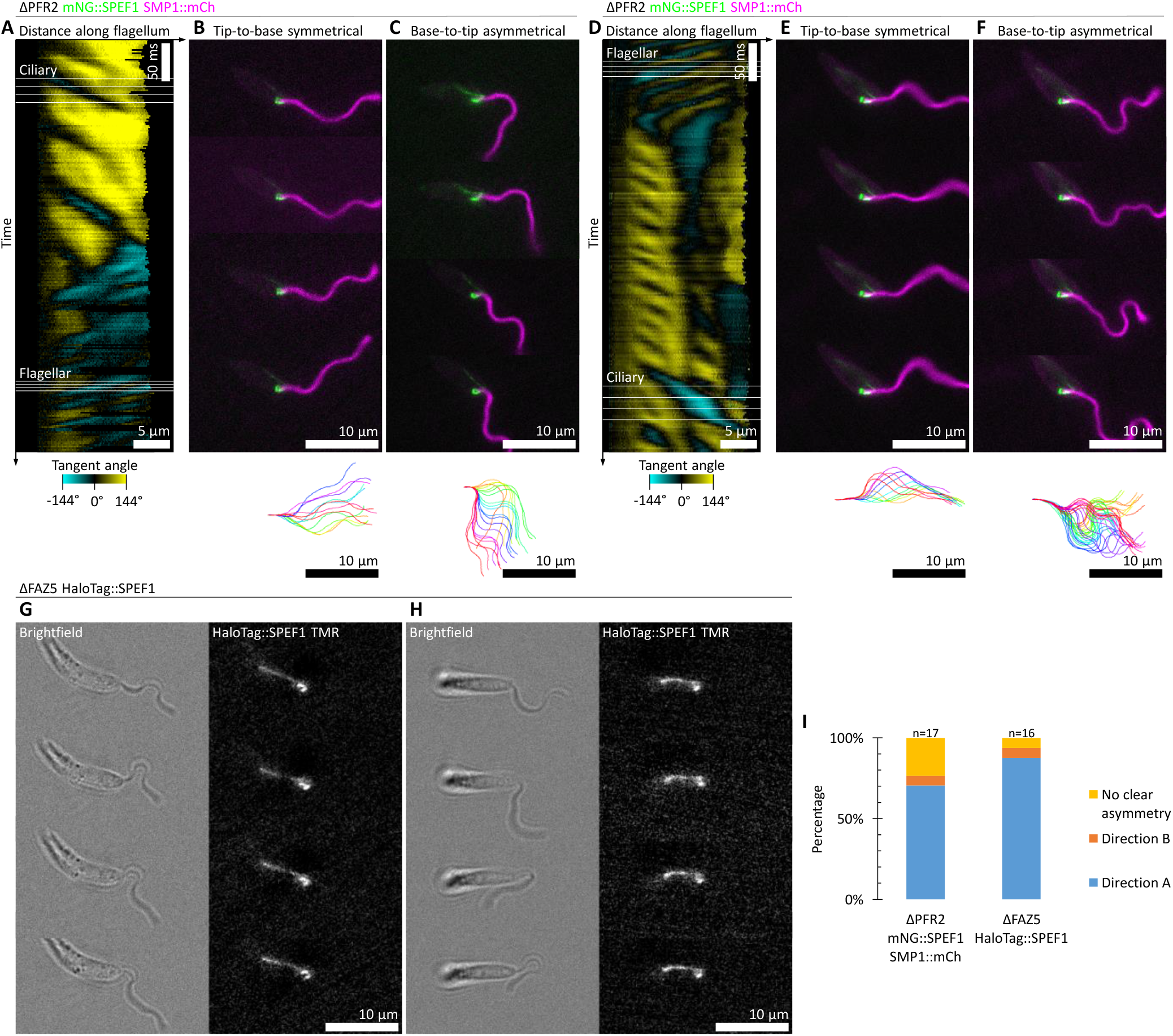
Neither disruption of the paraflagellar rod or the flagellum attachment zone inverts asymmetric beat polarity. **A-F.** Flagellum movement in two example cells of a PFR2, a paraflagellar rod protein, deletion mutant expressing mNG::PF16 and mCh::RSP4/6 (two axonemal proteins –central pair and radial spokes respectively) switching from a symmetric tip-to-base to an asymmetric base-to-tip beat, derived from a 100 Hz high-frame rate dual colour epifluorescence video. **A.** Flagellum curvature over time, as in Figure 3C, automatically traced from mNG::PF16 signal. **B.** Four frames showing the symmetric tip-to-base beat and the flagellum configuration over the beat cycle. **C.** Four frames showing the asymmetric base-to-tip beat and the flagellum configuration over the beat cycle. **D-F.** As for A-C except for the second example cell with a base-to-tip beat with no clear asymmetry. **G-H.** Flagellum movement in two example cells of a FAZ5, a flagellum attachment zone protein, deletion mutant expressing HaloTag::SPEF1 and labelled with tetramethylrhodamine ligand. Both examples persistently underwent an asymmetric base-to-tip beat, summarised with four frames from a 100 Hz high-frame rate green light brightfield/red fluorescence video. **G.** Instances of asymmetric bending in the A or B direction (as defined in Figure 5), counted from high frame rate fluorescence videos of either PFR2 or FAZ5 deletion mutants. The asymmetric beat retains a strong preference for the power stroke to bend away from the side of the cell with the lysosomal microtubule, with more examples of reverse beats with unclear asymmetry in the PFR2 deletion.

Finally, we considered the asymmetrically-positioned flagellum attachment zone at the base of the flagellum. Previous work identified the protein FAZ5 as vital for any lateral attachment between the flagellum and the flagellar pocket neck, in turn giving rise to a motility defect (Sunter et al., 2019). To determine if this defect was associated with altered asymmetry of the base-to-tip beat we generated a FAZ5 deletion cell line expressing HaloTag::SPEF1, where HaloTag is a self-labelling protein tag which covalently binds a chloroalkane with an amine-linked fluorophore – in this case tetramethylrhodamine. HaloTag was used as the ΔFAZ5 cell line was, for an unknown reason, refractory to tagging of SPEF1 with conventional beta barrel fluorophores (mNG, eYFP, mCh). We were also unsuccessful at generating the ΔFAZ5 cell line using a single selectable marker, leaving insufficient selectable markers to also tag SMP1. Therefore the cell line was visualised using a combination of red fluorescence and bright field transmitted light using a green filter for simultaneous visualisation of the flagellum and the orientation of the cell using HaloTag::SPEF1 (Figure 7G,H, Video 5). Flagella almost all underwent a base-to-tip beat, suggesting a reduced ability for a tip-to-base beat is the origin of the previously observed swimming defect (Sunter et al., 2019). Intriguingly, this suggests the flagellum attachment zone represses the base-to-tip beat. The bright field videos had insufficient contrast for automated tracing of beat curvature, but the base-to-tip beats appeared normal and consistently occurred with normal polarisation (Figure 7G,H, Video 5). The combined evidence from the ΔPFR2 (*n* = 17) and ΔFAZ5 (*n* = 16) mutants suggest the most prominent asymmetric flagellum-associated structures in *Leishmania* are not required for achieving asymmetric static flagellum curvatures (Figure 7I, cf. Figure 3,5).

## Discussion

*Leishmania* are highly genetically tractable cells (Beneke et al., 2017) and the promastigote life cycle stages have a canonical 9+2 single flagellum which switches between a near-planar (Walker and Wheeler, 2019) symmetric tip-to-base beat and an asymmetric base-to-tip (Gadelha et al., 2007; Holwill and McGregor, 1975) making it an excellent system for understanding the fundamental biology of flagella/cilia and control of their flagellar beating. In particular, new opportunities to analyse the origin and switching of beat asymmetry arise as the symmetric beat propagates from tip-to-base and the asymmetric beat from base-to-tip.

*Leishmania* have a near-axially symmetric cell shape which makes it challenging to analyse cell orientation from transmitted light micrographs. To overcome this limitation, we developed the first dual colour high resolution and high frame rate fluorescence visualisation of live swimming cells and used this to visualise proteins endogenously tagged with genetically encoded fluorophores (mCh, mNG) or fluorophore-binding proteins (HaloTag), using either simultaneous capture of red and green fluorescence or red fluorescence along with green transmitted light. This allowed us to analyse polarisation of flagellum beating using asymmetric intracellular structures (Figure 3), despite the outward axially-symmetric appearance of *Leishmania.*

Most organisms appear to have a fixed polarisation of the asymmetric beat, however in a uniflagellate cell it may be advantageous for both polarisations of the asymmetric beat to occur to rotate the cell either right or left. Our analysis showed that the asymmetric *Leishmania* beat has a fixed polarisation (Figure 5) arising from a static curvature in the flagellum (Figure 3) which always occurs with the same polarisation with the PFR on the inside of the curve. As asymmetric *Leishmania* beats rotate the cell this effectively limits the cell to rotate in just one direction, constraining the possible chemotaxis or rheotaxis responses *Leishmania* can undergo. This suggests they require a run-and-tumble-like chemotaxis mechanism – however our analyses were of cells trapped in a thin space between a slide and a coverslip and with three degrees of freedom swimming behaviours may differ somewhat.

In *Chlamydomonas,* regulation of the symmetric dynamic curvature of the flagellum to make a wave propagate and the static curvature of the flagellum to introduce asymmetry are separable, and some mutants (such as *mbo2*) cannot form a static curvature giving an aberrant symmetric waveform (Brokaw and Kamiya, 1987; Geyer et al., 2016; Segal et al., 1984). In *Leishmania*, no mutants are yet known which lose the asymmetry of their base-to-tip beat – deletions of the outer dynein arm-associated proteins dDC2 and LC4-like do promote asymmetric or symmetric beats respectively, but these beats occur in their normal base-to-tip and tip-to-base propagation directions respectively (Edwards et al., 2018). However, *Leishmania* deletion mutants (including PF16, Hydin, IC140) are known where the flagellum is paralysed but often still has a static curvature (Beneke et al., 2019) while, naïvely, paralysed flagella would be expected to be straight. Deletions of IC140 and Hydin are the most dramatic examples. The static curvature in these mutants is still polarised (Figure 6), however this polarisation is in the opposite direction to base-to-tip asymmetric beats (Figure 5). This strongly suggests flagellar polarisation is retained in the absence of the central pair complex and in the absence of the IC140-associated inner dynein arms, although the precise correspondence of this mutant phenotype to normal symmetric or asymmetric beats is unclear.

*Leishmania* has additional asymmetric flagellum associated structures which may have been important for conferring flagellum polarisation. However, loss of lateral flagellum attachment by the FAZ, which is analogous to rootlet structures in other organisms, did not reduce asymmetry or invert the polarisation of the asymmetric base-to-tip beat (Figure 7). Similarly, loss of almost the entire bulk of the PFR, a large and complex extra-axonemal structure with multiple potential functions (Portman and Gull, 2010), did not invert the polarisation of the asymmetric base-to-tip beat (Figure 7). While the FAZ and PFR are specific to the *Leishmania* branch of life, it suggests that asymmetric extra-axonemal structures and rootlet structures in other species are not responsible for the polarisation of asymmetric flagellum beats.

Together, this work greatly constrains the molecular origin of asymmetry in *Leishmania* base-to-tip beats. Our previous work indicated outer dynein arm-associated factors are likely important for switching between symmetric tip-to-base and asymmetric base-to-tip beats (Edwards et al., 2018), however controlling switching is distinct from the actual generation of asymmetry. The most likely remaining candidate is differences between the outer doublet decorations, in particular in the region of inner arm dynein b based on cryo-electron tomography of *T. brucei* (Imhof et al., 2019). However, the mechanical properties of the nine-fold asymmetric outer microtubule doublets of the axoneme itself could also be important. Specialised inner dynein arm are also implicated in *Chlamydomonas* flagellum movement asymmetries (Bui et al., 2009), perhaps indicating that this is the eukaryote-wide origin for asymmetry of flagellum movement.

## Methods

Procyclic promastigote *L. mexicana* expressing Cas9 and T7 polymerase, derived from WHO strain MNYC/BZ/62/M379 (Beneke et al., 2017) were grown in M199 medium with Earle’s salts and L-glutamine (Life technologies) supplemented with 26 mM NaHCO_3_, 5 μg/ml haemin, 40 mM, HEPES·NaOH (pH 7.4) and 10% FCS. *L. mexicana* were grown at 28°C and maintained at culture densities between 1×10^5^ to 1.0×10^7^ cells/ml.

For endogenous tagging of *L. mexicana* genes, constructs and sgRNAs were generated using the PCR-based approaches previously described (Beneke et al., 2017; Dean et al., 2015), using the pLPOT (also called pLrPOT) (Edwards et al., 2018) series of plasmids as the PCR template. *L. mexicana* were transfected and subjected to drug selection as previously described (Dean et al., 2015). For endogenous tagging with HaloTag, a new pLPOT variant with HaloTag and blasticidin deaminase was generated.

The *L. mexicana* cell lines with deletion of both alleles of flagellum and FAZ proteins were generated using the PCR-based approach previously described, using the pT series of plasmids as the PCR template (Beneke et al., 2017). All deletion cell lines have been previously been characterised: ΔPFR2, ΔIC140 and ΔHydin in Beneke et al., 2019 and ΔFAZ5 in Sunter et al., 2019.

Microscopy was performed with an Axio Observer A1 (Zeiss) microscope with incubator chamber using a 63× NA 1.4 Ph3 objective or a 100× NA 1.4 without phase ring using a 120 V metal halide fluorescence light sources (Zeiss, HXP 120 V). Standard fluorescent microscopy was performed with an mRFP (Zeiss, 63HE) or GFP (ThorLabs, MDF-GFP2) filter cube.

To synchronously record red and green fluorescence at high frame rate we used an OptoSplit II (Cairn Research) optical splitter. A custom microscope filter cube was fitted with a dual band (green and red) dichroic (Chroma Technology, 59022bs) and a dual band (blue and yellow) excitation filter (Chroma Technology, 59022x). The optical splitter was fitted with a filter cube with a 565 nm dichroic filter (Chroma Technology, T565lpxr) and green reflected and red transmitted light filters (Chroma Technology, ET520/40m and ET632/60m). When using white epi-illumination this results in the splitter projecting a green and red fluorescence image onto two halves of a single camera, a Neo 5.5 (Andor). A 2019 mm focal length lens was used in the red light path for focus correction.

To synchronously record red fluorescence and brightfield microscopy (in green) the sample did not emit green fluorescence so when using white epi-illumination and trans-illumination through a green filter (ThorLabs, Astronomy Green) this resulted in the splitter projecting a green brightfield and red fluorescence image onto the two halves of the camera.

The red and green images must be aligned to generate composite images. Images for calibration of position and scale/magnification were captured using multi-wavelength fluorescent beads (TetraSpeck 0.1 μm Microspheres, Invitrogen T7279), then the red and green images aligned using the same approach we previously used for multi-focal plane microscopy (Walker and Wheeler, 2019).

For microscopy of cells adhered to glass, a sample of cells from late logarithmic growth (0.5×10^7^ to 1.0×10^7^ cells/ml) were taken from culture, washed and placed on a slide then imaged live as previously described (Halliday et al., 2018).

For microscopy of free-swimming and naturally behaving cells samples were taken from cultures in late logarithmic growth. To reduce cell adherence, glass slides were blocked with BSA prior to use by immersion in 1% BSA in distilled water for 30 sec, followed by three washes in distilled water and air drying. 1 μl cells from culture in normal culture medium were applied to a 2 by 1 cm rectangle marked on a slide and coverslip using a hydrophobic pen, resulting in a liquid layer 2 μm thick.

In order to generate the plots of tangent angle at different distances along the flagellum over time, flagella were automatically traced from videomicrographs using ImageJ. We used the intensity thresholding, skeletonisation and tracing scheme we previously developed for phase contrast videos (Walker and Wheeler, 2019). For tracing the fluorescence high speed videos the approach was adapted such that red fluorescent signal (either SMP1::mCh or mCh::RSP4/6) was subject to intensity thresholding following a correction for photobleaching. Flagella were digitally straightened using the ImageJ straighten tool using the flagellum midline from the red fluorescence as the line selection for straightening both the green and red fluorescence images.

For SEM cytoskeleton preparations, cells were harvested by culture by centrifugation, washed three times in PBS, settled on coverslips in 24 well plates, washed with 0.1% NP-40 in PBS, washed three times in PBS and then fixed in 2.5% glutaraldehyde in PBS for 2 hours at room temperature and then at 4°C overnight. Fixed cells were washed three times for 5 min in PBS and incubated in 1% osmium tetroxide in PBS at 4°C for 1 hour in darkness. Samples were then washed three times with water for 5 min and ethanol dehydrated. Sample drying was completed by critical point drying using an Autosamdri-815 (Tousimis). Coverslips were then sputter gold coated for 60 s. Samples were imaged at around 20,000 to 35,000× on a JSM-6390 SEM (JEOL) with an Everhart-Thornley secondary electron detector using an EHT target of 2 kV and a 20 μm aperture.

## Acknowledgements

I would like to thank Jack Sunter for kindly providing the FAZ5 deletion mutant.

RJW and ZW were supported by the Wellcome Trust (211075/Z/18/Z, 104627/Z/14/Z). EG was supported by the Royal Society (UF100435 and UF160661). TB was supported by the MRC (15/16_MSD_836338).

## Author contributions

ZW and TB generated the cell lines. TB performed the scanning electron microscopy. High framerate fluorescence light microscopy and data analysis was developed and performed by RJW. RJW wrote the manuscript and all authors contributed to data analysis and manuscript revision.

---

**Figure S1.**
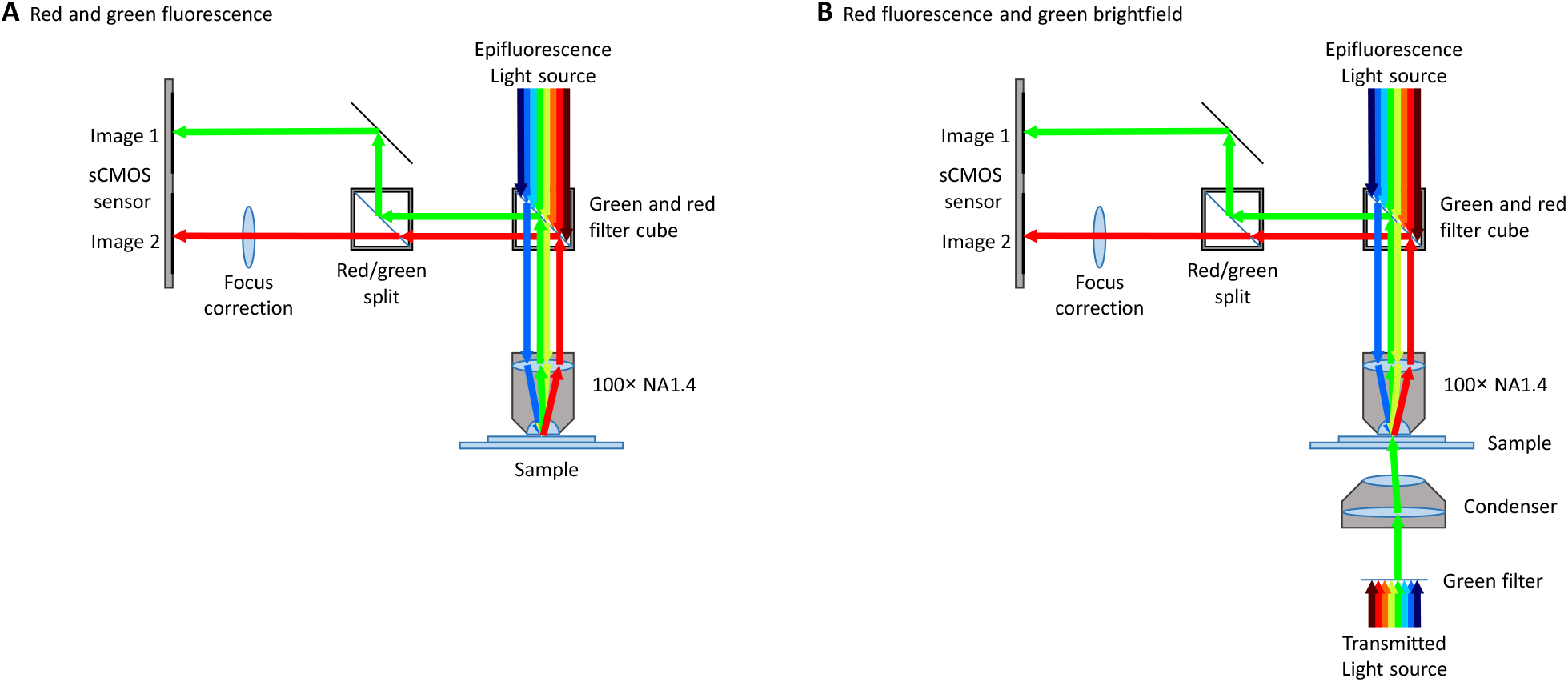
Summary of the microscope light paths for dual colour high resolution high frame rate microscopy. **A.** The light path configuration for simultaneous green and red fluorescence visualisation. **B.** The light path for red fluorescence visualised simultaneously with bright field transmitted light. The light path is the same, except that the sample does not emit green fluorescence and a transmitted light source through a green filter is used.

Video 1. Animated version of Figure 2.

Video 2. Animated version of Figure 3.

Video 3. Animated version of Figure 4.

Video 4. Animated version of Figure 7A-F.

Video 5. Animated version of Figure 7G-H.

